## Supplemental Figures 1 - 9 for "PIKfyve controls dendritic cell function and tumor immunity"

Extended Data Figure 1

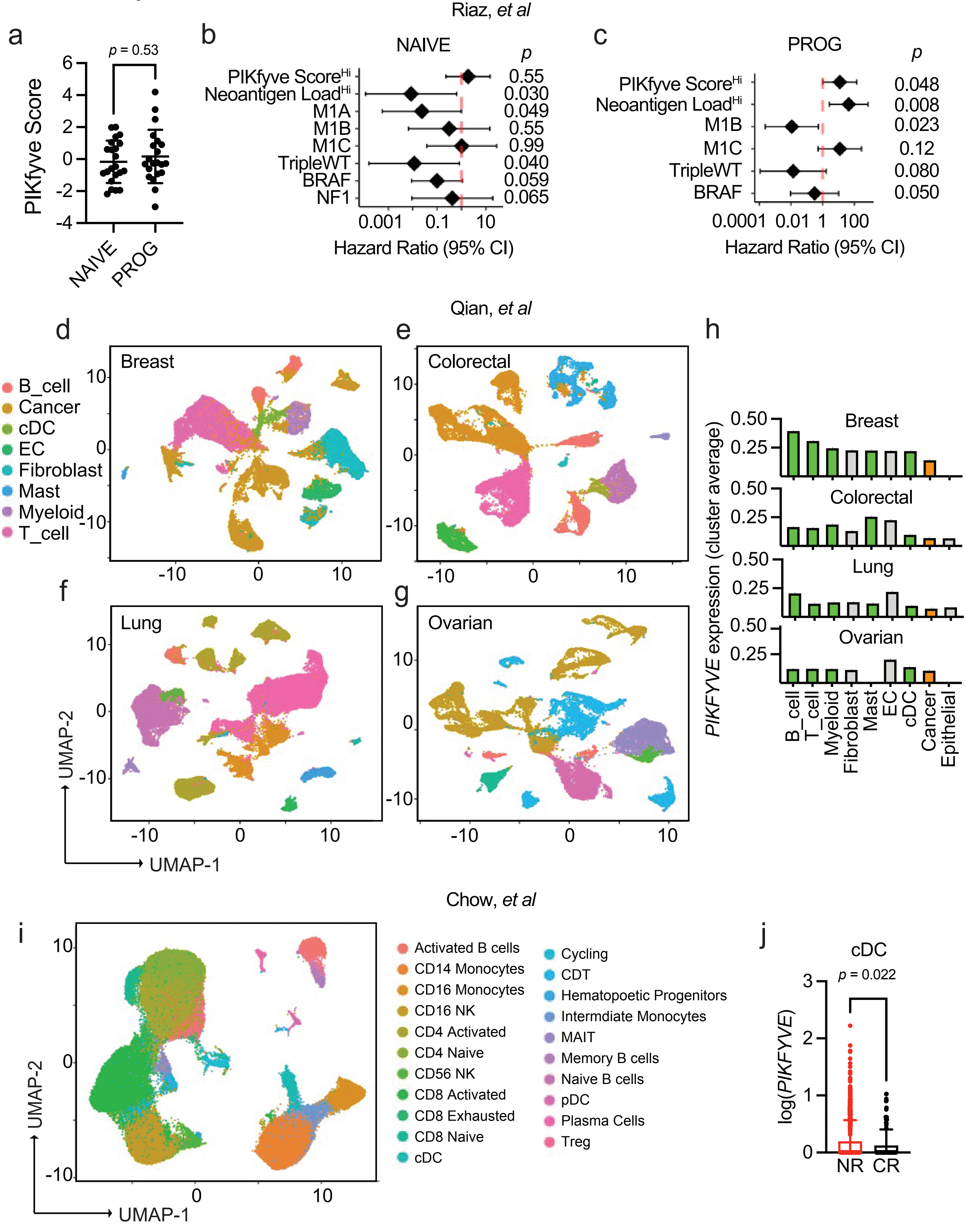

Extended Data Figure 2

a

Gating strategy for Supp 2b-g

| Subset | Gating Strategy |
|---|---|
| Naïve | $CD3^{+}CD8^{+}CD44^{Lo}CD62L^{+}$ |
| Terminally differentiated | $CD3^{+}CD8^{+}CD44^{Hi}$KLRG1+ |
| Resident Memory | $CD3^{+}CD8^{+}CD44^{Hi}KLRG1^{-}$CD49a+ |
| Exhausted | $CD3^{+}CD8^{+}CD44^{Hi}$KLRG1-CD49a-$TIM3^{+}PD\text{-}1^{+}$ |
| Central memory | $CD3^{+}CD8^{+}CD44^{Hi}$KLRG1-CD49a-$TIM3^{-}PD\text{-}1^{-}CD62L^{+}$ |
| $CD69^{+}$ Effector | $CD3^{+}CD8^{+}CD44^{Hi}$KLRG1-CD49a-$TIM3^{-}PD\text{-}1^{-}CD62L^{-}CD69^{+}$ |
| $CD69^{-}$ Effector | $CD3^{+}CD8^{+}CD44^{Hi}$KLRG1-CD49a-$TIM3^{-}PD\text{-}1^{-}CD62L^{-}CD69^{-}$ |

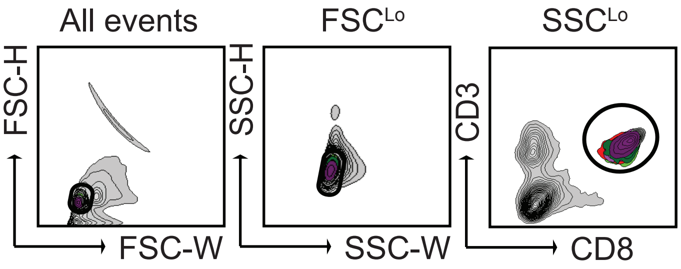


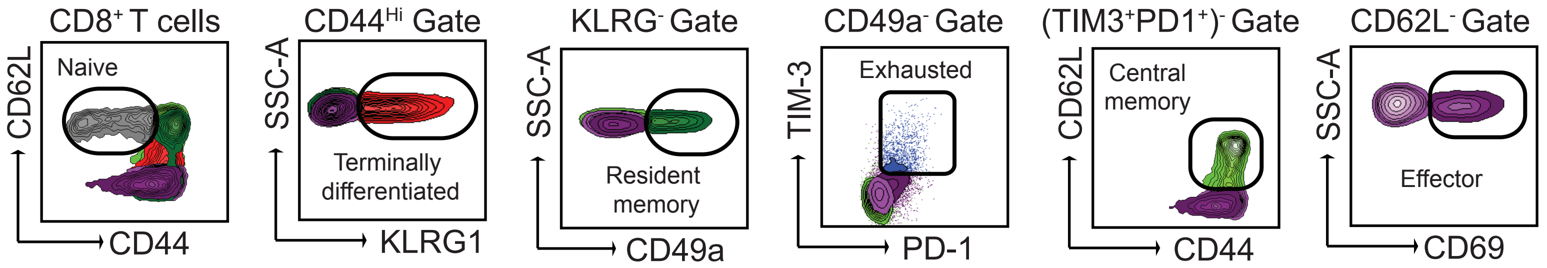


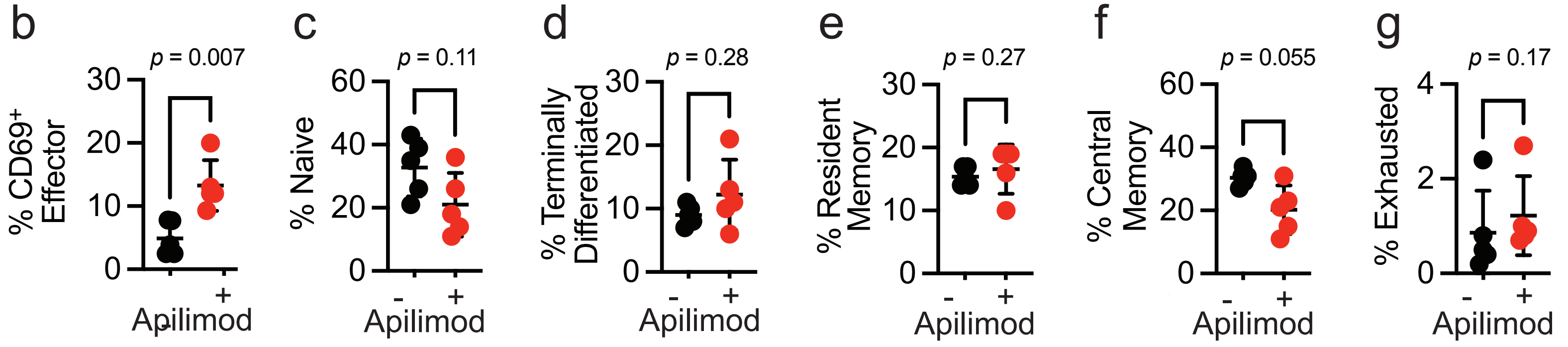


h

Gating strategy for Fig.2a-b, Supp 2i-l

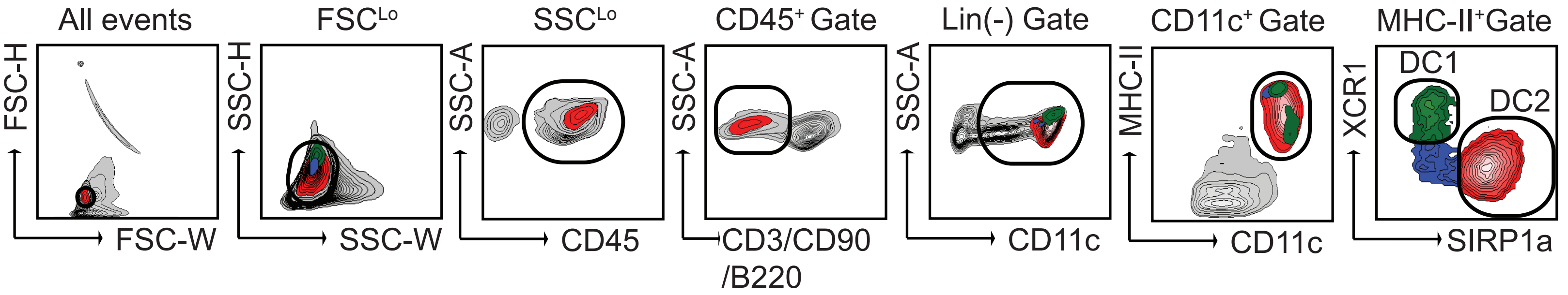


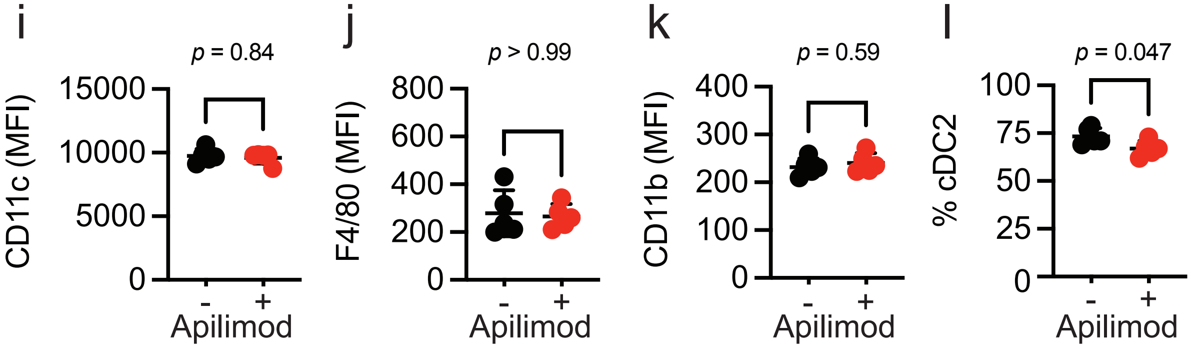

Extended Data Figure 3

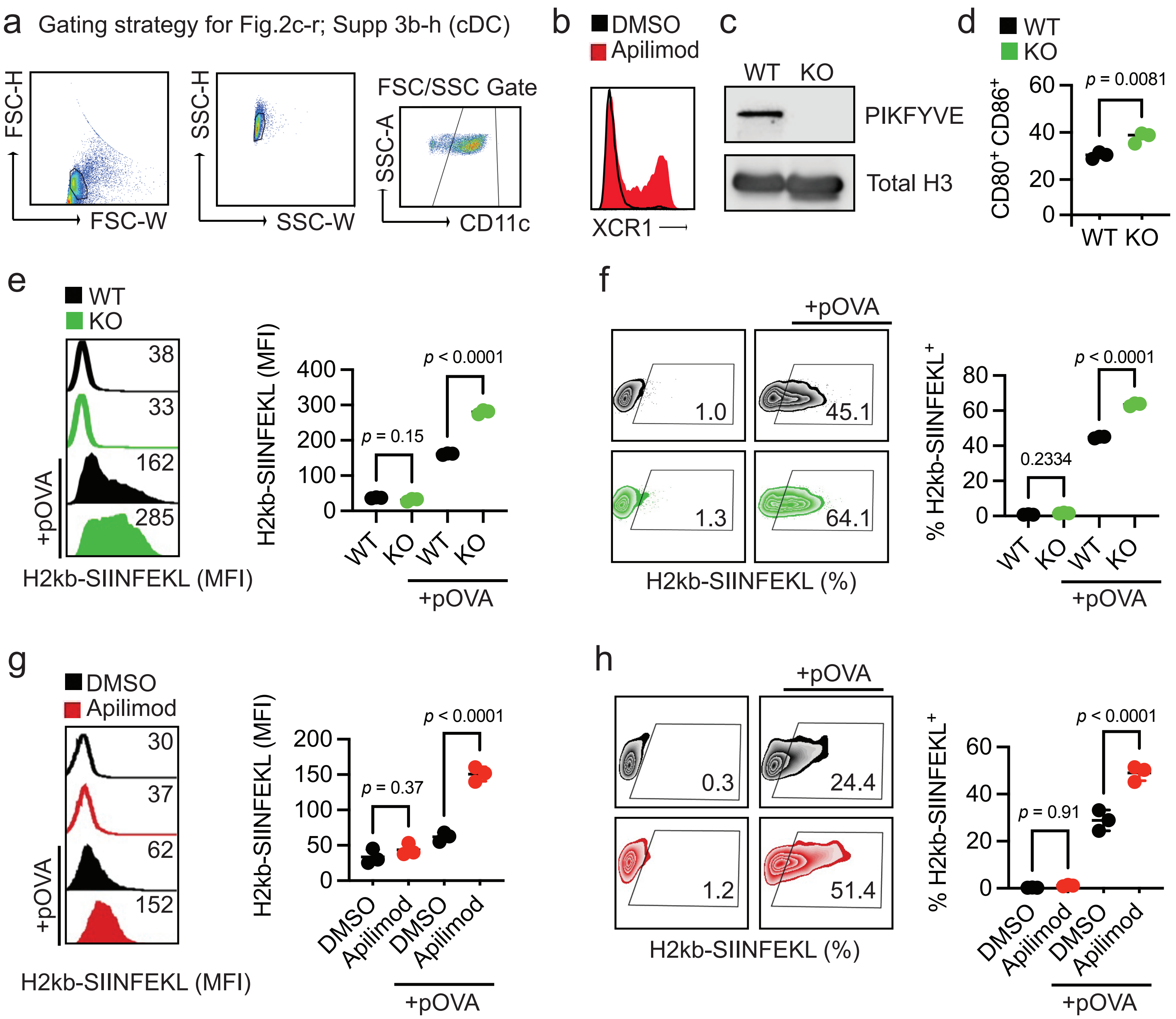

Extended Data Figure 4

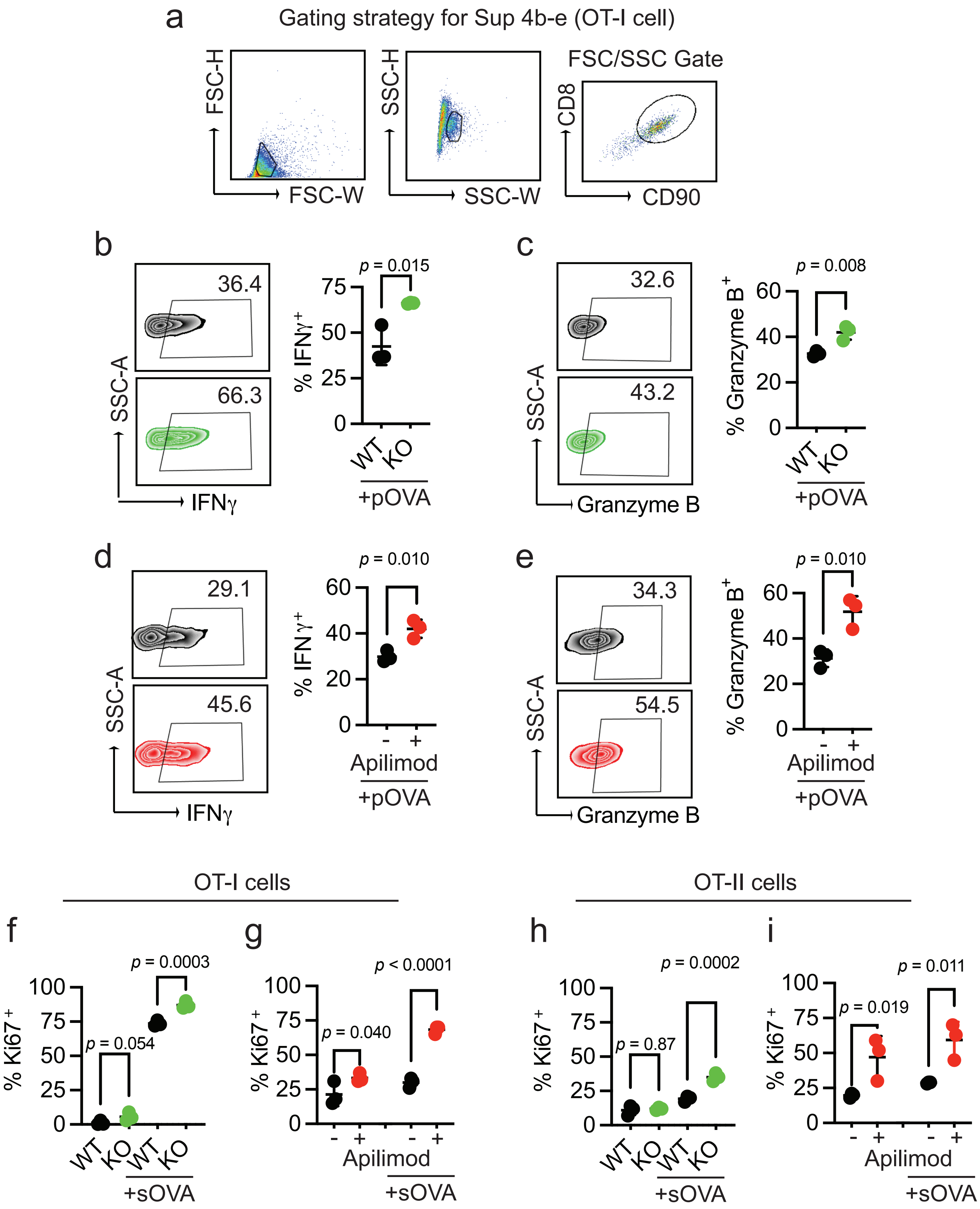

Extended Data Figure 5

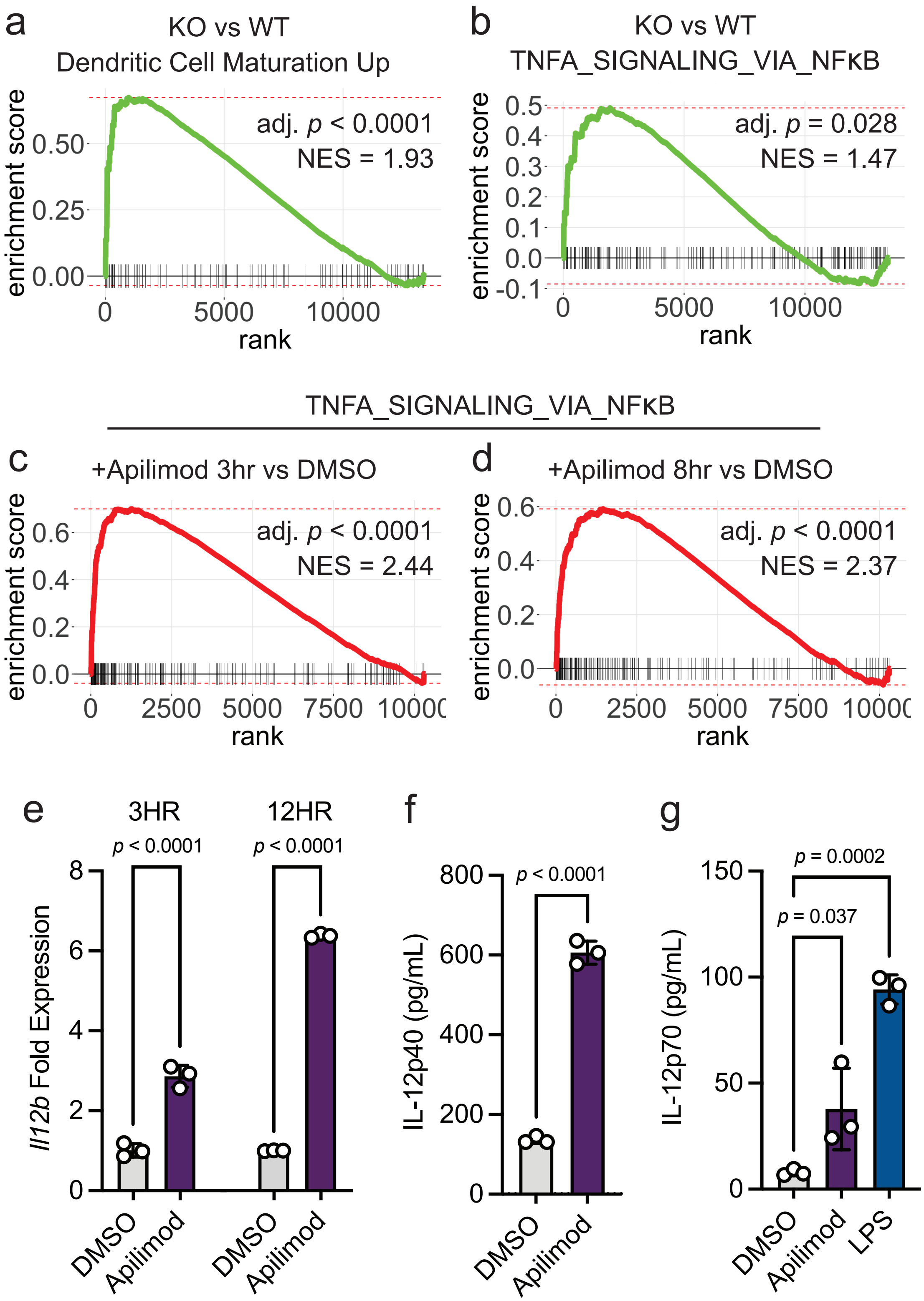

Extended Data Figure 6

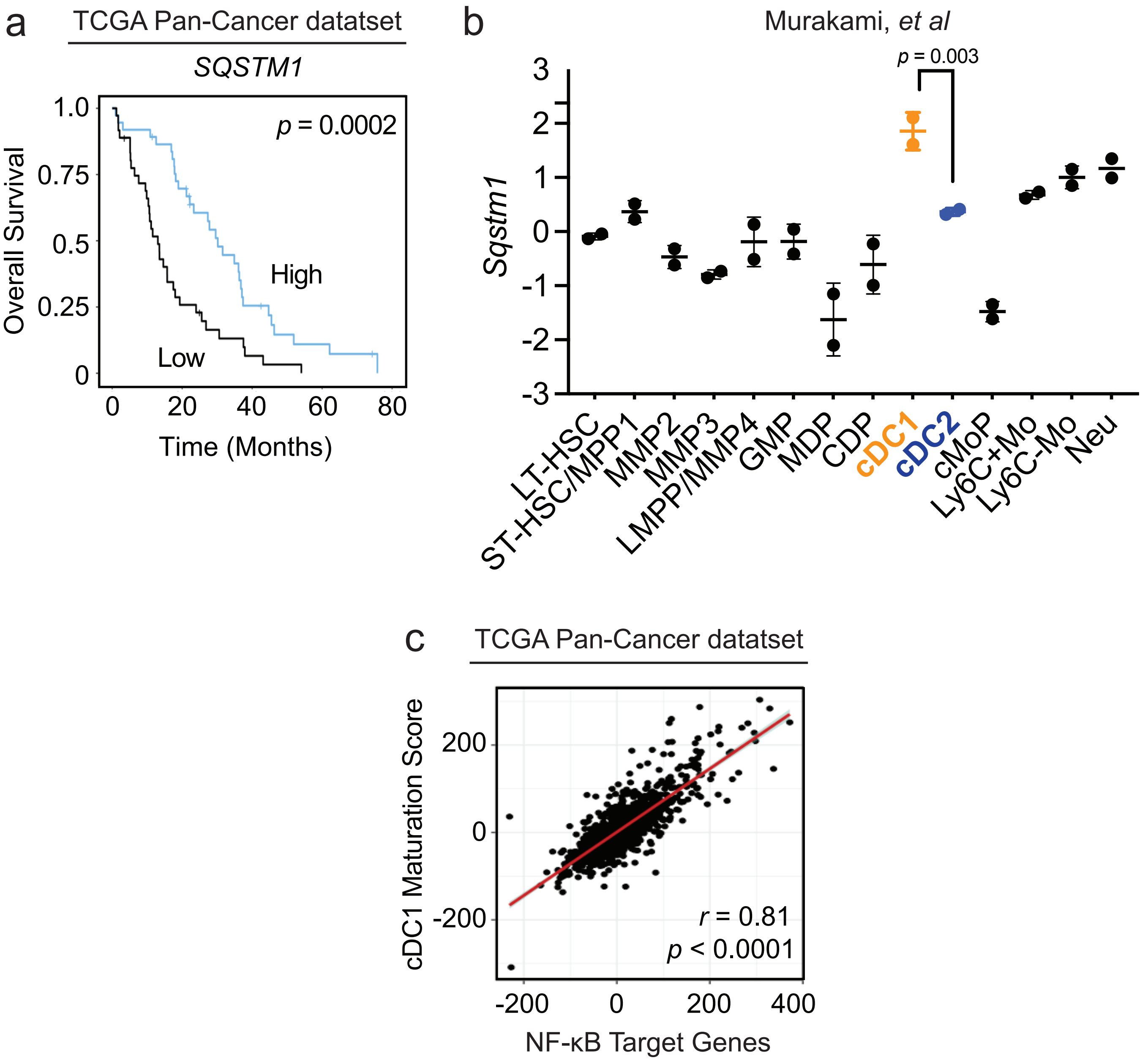

Extended Data Figure 7

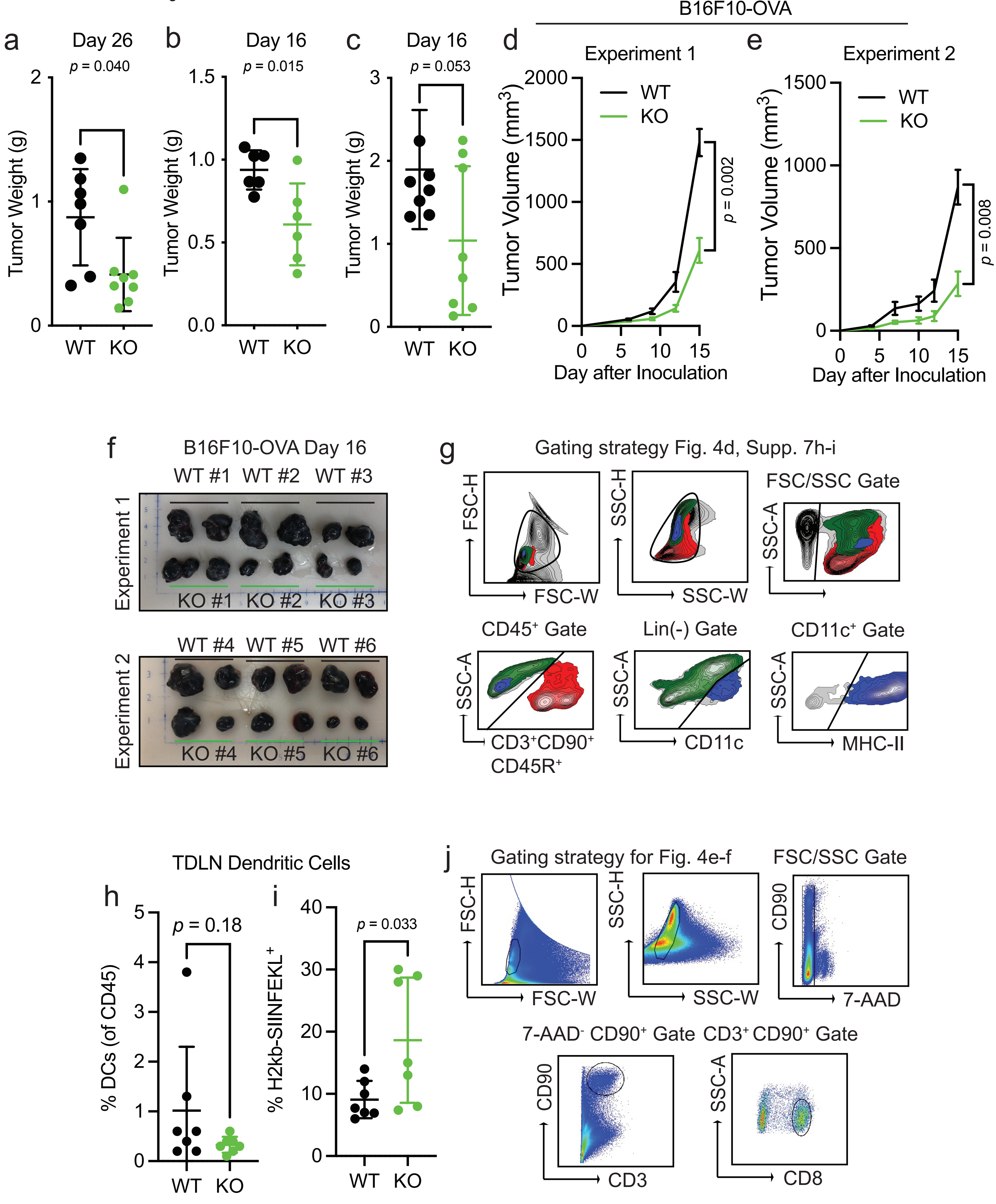

Extended Data Figure 8

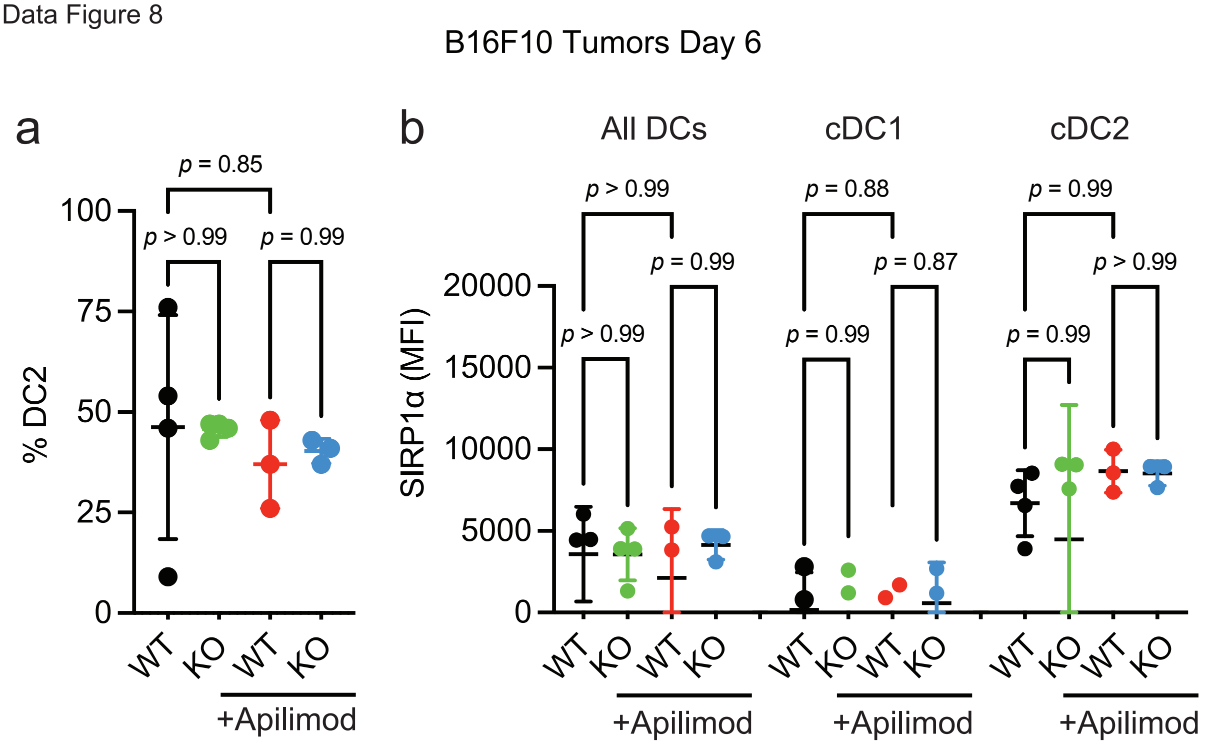

Extended Data Figure 9

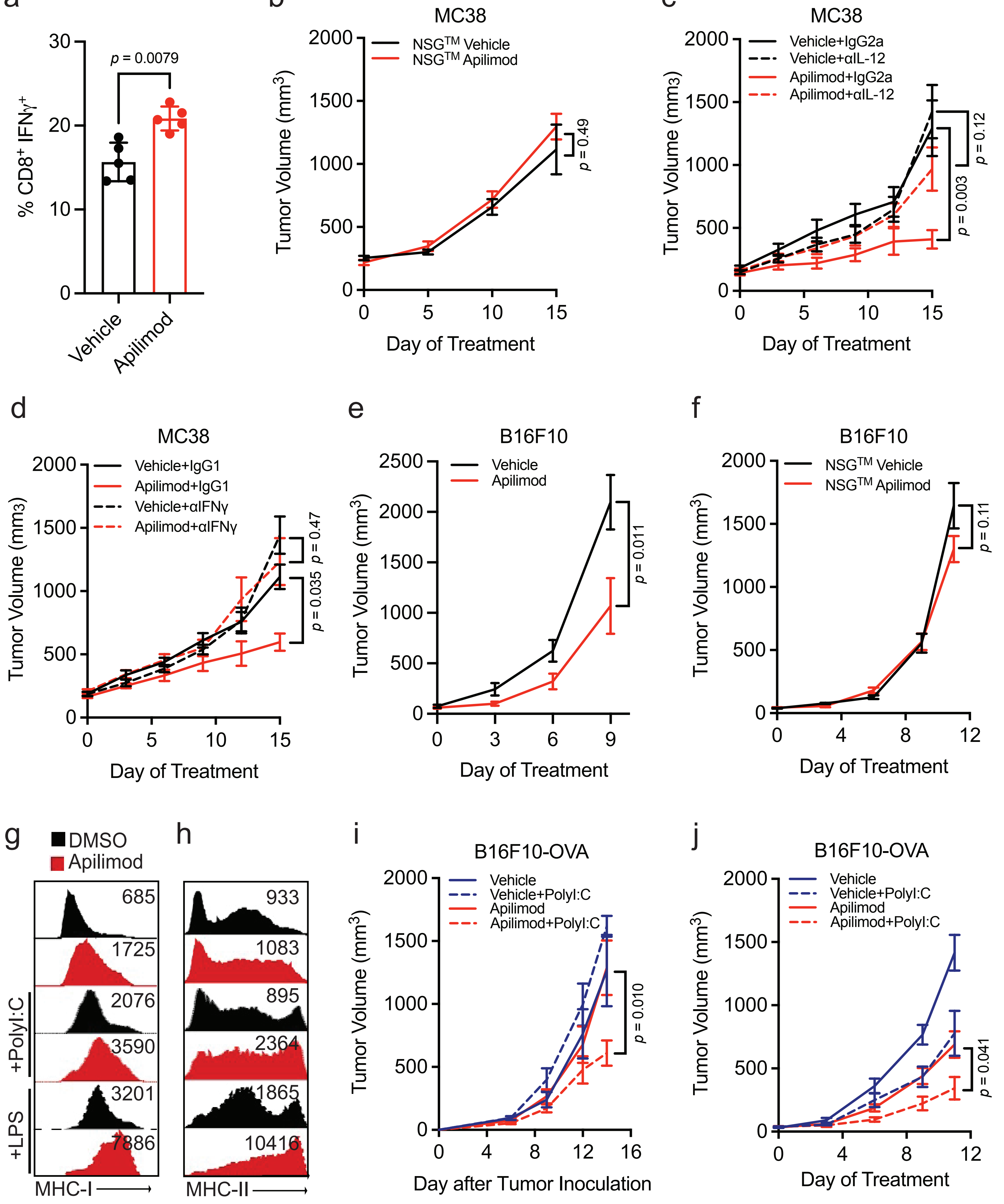
